# Australian rodents reveal conserved Cranial Evolutionary Allometry across 10 million years of murid evolution

**DOI:** 10.1101/2020.04.30.071308

**Authors:** Ariel E. Marcy, Thomas Guillerme, Emma Sherratt, Kevin C. Rowe, Matthew J. Phillips, Vera Weisbecker

## Abstract

Among vertebrates, placental mammals are particularly variable in the covariance between their cranial shapes and body size (allometry), with the notable exception of rodents. Australian murid rodents present an opportunity to assess the cause of this anomaly because they radiated on an ecologically diverse continent unique for lacking other terrestrial placentals. Here we used 3D geometric morphometrics to quantify species-level and evolutionary allometries in 38 species (317 crania) from all Australian murid genera. We ask if ecological opportunity resulted in greater allometric diversity; conversely, we test if intrinsic constraints and/or stabilizing selection conserved allometry. To increase confidence in species-level allometric slopes, we introduce a new phylogeny-based method of bootstrapping and randomly resampling across the whole sample. We found exceedingly conserved allometry across the 10 million year split between *Mus* and the clade containing Australian murids. Cranial shapes followed craniofacial evolutionary allometry (CREA) patterns, with larger species having relatively longer snouts and smaller braincases. CREA is consistent with both intrinsic constraints and stabilizing selection hypotheses for conserved allometry. However, large-bodied frugivores evolved faster, while carnivorous specialists showed skull modifications known to conflict with masticatory efficiency. These results suggest a strong role of stabilizing selection on the masticatory apparatus of murid rodents.

## Introduction

Allometry, or the scaling relationships between physical traits as body size changes, is a pervasive pattern in the evolution of animal morphological diversity (Huxley and Teissier 1936). Related species with different body sizes usually have morphologies close to those predicted by their clade’s evolutionary allometric trajectory, even when natural selection would favor alternative scaling relationships (Pélabon et al. 2014; Serb et al. 2017). Therefore, evolutionary allometry represents a compromise between the natural selective regimes driving diversification and the clade’s inherited development underlying morphology (Voje et al. 2014). Placental mammals show exceptional variation in size and morphology and thus offer an intriguing case to explore this compromise between extrinsic selection and intrinsic development (Tsuboi et al. 2018). Indeed, the unique placental pregnancy appears to provide a developmental environment that increases the viability of early developmental variations compared to other vertebrates, including other mammals (Lillegraven 1974; Millar 1977). In turn, greater allometric diversity provides natural selection with more morphological diversity to target, which could facilitate both rapid allometric divergence (Esquerré et al. 2017) and increased speciation in placentals (Jungers 1982; Schluter 1996; Wund et al. 2012; Marcy et al. 2016).

Rodents deviate from the correlation between species richness and morphological diversity observed in other placental mammals (Hautier and Cox 2015). Muridae, a single rodent family, includes 12.8% of all mammalian species but their morphology appears to follow a highly conserved allometric pattern (Fabre et al. 2012; Burgin et al. 2018), especially within the cranium (Firmat et al. 2014; Verde Arregoitia et al. 2017; Alhajeri and Steppan 2018). The unexpected allometric conservatism of murid rodents positions them as model organisms for understanding how the interaction of extrinsic and intrinsic factors impacts allometric variation and subsequent macroevolutionary patterns.

The relative importance of extrinsic natural selection and intrinsic developmental processes on allometric patterns has long been debated (Pélabon et al. 2014). Their relative importance likely exists along a spectrum, but there are three main hypotheses that attempt to define distinct, testable categories (Brigandt 2015). The first hypothesis – which most placental mammals seem to illustrate – posits that disruptive (or directional) selection can alter allometric patterns quickly, especially when a new selective pressure emerges (Frankino et al. 2005; Tsuboi et al. 2018). This “extrinsic pressure hypothesis” expects changes in selection to be the most important determinant of the allometric patterning for a given species or clade. At the opposite end of the spectrum, the second hypothesis emphasizes how conserved allometric patterns arise from inherited developmental processes (Voje et al. 2014). This “intrinsic constraint hypothesis” posits that allometry stays conserved because genetic changes to development have pleiotropic effects and thus expects allometry to be limited to the few viable variations (Marroig and Cheverud 2010; Shirai and Marroig 2010). The intermediate hypothesis posits that the interaction of extrinsic stabilizing selection (a subcategory of natural selection) on intrinsic development produces consistently functional morphologies (Marroig and Cheverud 2005). Unlike the extrinsic pressure hypothesis, this “stabilizing selection hypothesis” expects outcomes similar to – perhaps even indistinguishable from—the intrinsic constraint hypothesis (Brigandt 2015). Notably, the stabilizing selection hypothesis expects that sustained stabilizing selection on limited viable genetic variation could hone an allometric trajectory that facilitates a clade’s rapid radiation (Marroig and Cheverud 2005; Voje et al. 2014; Cardini et al. 2015). This so-called “allometric line of least resistance” is thought to scale stable, functional morphological ratios for a wide range of body sizes (Schluter 1996).

The allometric patterning of Australian murid rodents could plausibly be characterized by each of the three hypotheses. First, their radiations would have experienced new extrinsic selection pressures by immigrating from wet tropics onto a much drier continent (Aplin and Ford 2014; Smissen and Rowe 2018). Indeed, unlike nearly all other murid radiations, the Australia-New Guinea radiations show some evidence of following an ecological opportunity model (*sensu* Yoder et al. 2010), where adaptation to new environments, especially the dry habitats, could be driving speciation (Schenk et al. 2013; Smissen and Rowe 2018; but see Alhajeri et al. 2016). Furthermore, Australia uniquely lacks other terrestrial placental mammals (Aplin and Ford 2014), therefore it is possible that a release from competition could allow extrinsic pressures to push murid rodent allometry into morphological niches unavailable to all other murids. However, in order for extrinsic pressures to be the main determinant of allometric patterns, murids would need to arrive in Australia with flexible developmental processes. Evidence for conserved allometry in murids in general (Porto et al. 2013; Firmat et al. 2014) makes the extrinsic pressure selection hypothesis appear unlikely for Australian murids.

Additional understanding on the intrinsic factors influencing allometry in mammals can come from assessments of allometry-related shape variation patterns. Many major mammalian clades have conserved shape patterns that follow the proposed “rule” for craniofacial evolutionary allometry (CREA; Cardini et al. 2015), where larger species have relatively longer snouts and smaller braincases (Radinsky 1985; Cardini and Polly 2013; Cardini et al. 2015; Tamagnini et al. 2017; Cardini 2019). CREA does not yet have a satisfactory explanation (Cardini 2019), but this conserved allometric pattern could be attributed to post-natal growth patterns instrinsic to both marsupial and placental mammals (Cardini et al. 2015). If CREA is present in rodents, it could also possibily be explained by the stabilizing selection hypothesis because cranial allometry could scale the function of their derived masticatory apparatus for gnawing (Alhajeri and Steppan 2018). The apparatus includes actively sharpened incisors, a diastema allowing independent occlusion at the incisors or at the molars, and a craniomandibular joint allowing movement between occlusion points (Druzinsky 2015). This complexity would decrease viability of developmental alterations since any maladaptive ratios would decrease fitness and simultaneously reinforce an allometric line of least resistance. However, many murid dietary specialists diverge in mandible shape (Renaud et al. 2007, 2007; Esselstyn et al. 2012; Fabre et al. 2017). These specialists indicate an interesting threshold between stabilizing and other forms of natural selection, which suggests the latter can shift long-standing allometric patterns to accommodate new masticatory biomechanics. Therefore, exceptions to the allometric “rules” may provide insight into the conditions leading to large adaptive leaps, such as a population entering a new selection regime, evolving a genetic mutation that lifts a constraint, or both (e.g. Polly 2008; Cardini et al. 2015).

Australian murid rodents represent at least eight recent and relatively rapid radiations with high species richness and diverse ecological adaptations, including dietary and locomotor specializations (Rowe et al. 2008; Aplin and Ford 2014). In this study, we use 3D geometric morphometric analyses to assess their cranial allometry and morphology within and among 38 species, covering 58% of species and all genera extant on modern-day Australia (fig. 1). Specifically, we ask three questions:

1) Are there divergent allometric patterns, consistent with the ecological opportunity model of a predominant role for extrinsic pressures on Australian murid rodent allometry?
2) If allometry is conserved, does it follow suggested deeply conserved mammalian shape patterns like CREA?
3) Furthermore, if allometry is conserved, is there evidence for stabilizing selection, in particular an “allometric line of least resistance” facilitating species to rapidly evolve functional shapes along the common evolutionary allometric trajectory?

**Figure 1:**
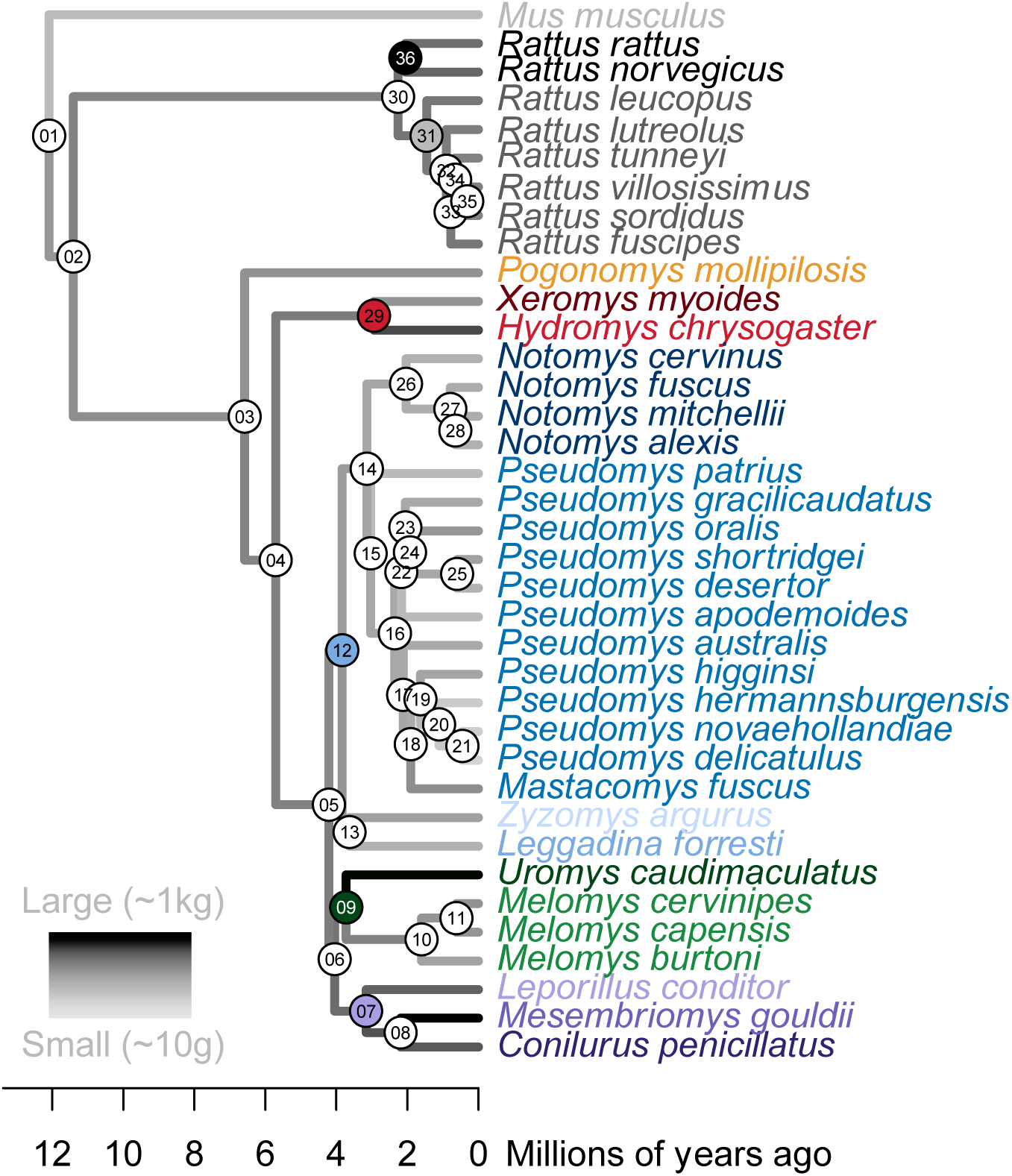
The time-calibrated molecular phylogeny generated for 37 of 38 species in this study. Node numbers correspond to those in figure 3. Filled nodes indicate the six major clades, whose ancestors each arrived to Australia at distinct times, after Aplin and Ford (2014). Species name colors were gradated across the genera in each major clade (e.g. blues for the Pseudomys division, node 12) and used consistently throughout. Phylogeny branches are tinted by body size (estimated from cranial centroid size (Zelditch et al. 2004)). These were generated by *phytools* (v.0.6- 99) function *plotBranchbyTrait* using species mean cranial centroid sizes, estimated for ancestors from these tips (Revell 2012).

## Methods

### Data Collection: Shape and Size Data

We sampled crania from four Australian museums: Queensland Museum (Brisbane), Australian Museum (Sydney), South Australian Museum (Adelaide), and Museums Victoria (Melbourne). The 317 adult specimens represent 35 species of native and 3 species of invasive rodents, including all 14 extant genera of rodents in Australia. Adults were determined by an emergent third molar and closure of the basisphenoid-basioccipital suture. When possible, species were represented by 10 individuals, 5 males and 5 females (see table A1). Each cranium was scanned with a HDI109 blue light surface scanner (LMI Technologies Inc., Vancouver, Canada) on a rotary table. We followed the same scanning method as Marcy et al. (2018). Note that our scanner’s resolution was insufficient to capture the very thin lateral zygomatic arches of smaller specimens, which we accepted as a trade-off for the large number of specimens acquired. This was deemed appropriate because skeletonization would have caused specimen preparation error as the fine structure dried and lost support from surrounding muscles (Yezerinac et al. 1992; Schmidt et al. 2010). The rest of the crania, including the roots of the zygomatic arches and main areas of muscle attachment (i.e. massetaric scar and temporal fossa), was captured.

3D crania scans were landmarked in Viewbox version 4.0 (dHAL software, Kifissia, Greece; www.dhal.com (Polychronis et al. 2013)). A preliminary analysis of all genera using the landmarking template from Marcy et al. (2018) identified the eastern chestnut mouse, *Pseudomys gracilicaudatus* (Gould, 1895) QM-JM9681 as the mean specimen, which was used to create a new template. In the present study, crania were characterized by 60 fixed landmarks, 141 curve semi-landmarks, and 124 patch semi-landmarks for a total of 325 landmarks (table A2 and fig. A1). The fixed landmarks do not slide, the curve semi-landmarks slide along a user-defined curve, and the patch semi-landmarks slide across a surface bounded by curves. Sliding was done in Viewbox by minimizing bending energy from 100% to 5% exponential energy over six cycles of projection and sliding.

During landmarking the mesh was rotated and/or the virtual lighting was changed to locate each landmarks’ position. The specimens were landmarked in a random order by one person (AEM) to avoid inter-observer error (Fruciano et al. 2017). The first 20 specimens were removed to reduce user error prior to learning the template. Another 20 specimens were digitized twice to assess observer error. Once landmarking was complete, large landmarking errors were identified and corrected with the *plotOutlier* function in *geomorph* (v.3.0.7) (Adams, Collyer, and Kaliontzopoulou 2018). Repeatability for the main dataset was about 93%, which is standard user error for 3D geometric morphometrics (e.g. Fruciano 2016; Fruciano et al. 2017; Marcy et al. 2018).

The landmark coordinates were prepared for statistical analysis using a generalized Procrustes analysis – removing differences in size, position, and orientation, leaving only shape variation (Rohlf and Slice 1990) – in R (v.3.6.1) (R Core Team 2019) and *geomorph* (v.3.1.0) (Adams, Collyer, and Kaliontzopoulou 2019). Afterwards, each cranium retains an associated centroid size as a proxy of body size (calculated as the square root of the sum of the squared distance of every landmark to the centroid or “center” of the landmark configuration (Zelditch et al. 2004)). The processed coordinates were used as shape variables for the following geometric morphometric, allometric, and phylogenetic analyses. While some reviews have criticized geometric morphometrics for using Gould-Mosimann allometry over the original Huxley-Jolicoeur framework (Pélabon et al. 2014; Voje et al. 2014), both frameworks are logically compatible and unlikely to yield contradictory results (Klingenberg 2016).

### Data Collection: Time-Calibrated Phylogenetic Tree

The phylogenetic tree (see fig. 1) for murid rodent species represented by 3D surface scans was compiled from DNA sequences from ten previously sequenced genes: a mitochondrial protein coding locus (cytochrome b) and 9 nuclear exons (exon 1 of ADRA2B, exon 9 of ARHGAP21, exon 11 of BRCA1, exon 8 of CB1, exon 10 of GHR, exon 1 of IRBP, the single exon of RAG1, exon 7 of TLR3, and exon 29 of vWF). Using the alignments of Smissen and Rowe (2018) as our starting point, we removed extraneous taxa and added taxa to obtain an alignment including 72 murid species in subfamily Murinae (table A3). These included all but two of the 38 species in our morphological dataset. No sequences were available for the central rock-rat, *Zyzomys pedunculatus* (Waite, 1896) or for the prehensile-tailed mouse, *Pogonomys mollipilosus* (Milne-Edwards, 1877). However, for our analyses we used the New Guinean large tree mouse, *Pogonomys loriae* (Thomas, 1897) as a surrogate for *P*. *mollipilosus* as the two species are equidistant from other taxa in our analyses. Additional species were included as outgroups and for fossil-calibration.

With our concatenated alignment of 10 loci and 72 species, we estimated a time-calibrated ultrametric phylogeny using a relaxed molecular clock approach in BEAST (v.2.1.3) (Bouckaert et al. 2014). Appropriate DNA sequence partitions and substitution models were found following settings as were a total of four calibration points specified in Smissen and Rowe (2018). These combine three fossils from the Siwalik formation (Kimura et al. 2015) with a calibration for the origin of Australian murines (Aplin and Ford 2014). We applied a Yule speciation prior and set the birthrate prior to exponential with an initial mean of 10. Other priors were left at default settings. Initial runs were used to optimize operators and we conducted a final Markov Chain Monte Carlo run with 2 × 10^8^ generations, sampling trees and other parameters every 2000 generations. We evaluated convergence and assessed sampling adequacy in Tracer (v.1.4) (Rambaut and Drummond 2007). TreeAnnotator was used to discard the first 20% of trees as burn-in and pool the remaining samples to form the posterior distribution and generate a maximum clade credibility tree. Finally, we manually pruned the resultant tree to the 37 species. This and other recent phylogenies show the broad-toothed rat, *Mastacomys fuscus* (Thomas, 1882) falling within genus *Pseudomys* (Smissen and Rowe 2018) so we grouped this species as part of *Pseudomys*.

### Allometric Variation

To address all three questions and characterize allometric patterns in Australian murids, we tested allometric variation at three levels: static allometry (species-level), evolutionary allometry (among clades), and a phylogenetic rarefaction testing every node in the tree.

First, variation in static (species-level) allometries was tested using an analysis of covariance (ANCOVA) model (see table 1*A*), implemented with *geomorph* function *procD*.*lm* (for highly-multivariate data), and evaluated for significance with Goddall’s (1991) F-test and 500 permutations. A post-hoc test using package *RRPP* (v.0.4.3) (Collyer and Adams 2018, 2019) function *pairwise* evaluated whether the static allometric slopes of all species (n = 38) significantly differ from one another (see table A4). Multiple comparisons were accounted for by reducing alpha to 0.01. The model was visualized by plotting the regression scores of shape on size versus log centroid size (see fig. 2) (Drake and Klingenberg 2008).

**Table 1:** ANCOVAs: Static allometry uses shapes and centroid sizes from all individuals from all 38 species (A). Evolutionary allometry uses mean shapes and mean centroid sizes of 37 species, which were then grouped into clades (fig. 1) (B). Abbreviations: degrees of freedom (df), sum of squares (SS), mean squares (MS), coefficient of determination (R_2_), F-statistic (F), effect size (Z), and P-value estimated from parametric F-distributions (Pr(>F)).

| A) Static Allometry (species-level) ANCOVA |  |  |  |  |  |  |  |
| --- | --- | --- | --- | --- | --- | --- | --- |
| | df | SS | MS | $R^2$ | F | Z | $\text{Pr}( > F )$ |
| Log(size) | 1 | 0.625 | 0.625 | 0.365 | 450. | 10.5 | 0.002 |
| Species | 37 | 0.694 | 0.019 | 0.405 | 13.5 | 19.9 | 0.002 |
| Log(size):species | 37 | 0.060 | 0.002 | 0.035 | 1.16 | 18.5 | 0.002 |
| Residuals | 241 | 0.335 | 0.001 | 0.195 |  |  |  |
| Total | 316 | 1.713 |  |  |  |  |  |
| B) Evolutionary Allometry (among clades) ANCOVA |  |  |  |  |  |  |  |
| | df | SS | MS | $R^2$ | F | Z | $\text{Pr}( > F )$ |
| Log(size) | 1 | 0.625 | 0.625 | 0.364 | 271 | 9.40 | 0.002 |
| Clade | 7 | 0.300 | 0.043 | 0.175 | 18.6 | 14.8 | 0.002 |
| Log(size):Clade | 7 | 0.096 | 0.014 | 0.056 | 5.94 | 10.8 | 0.002 |
| Residuals | 301 | 0.069 | 0.002 | 0.404 |  |  |  |
| Total | 316 | 1.71 |  |  |  |  |  |

**Figure 2:**
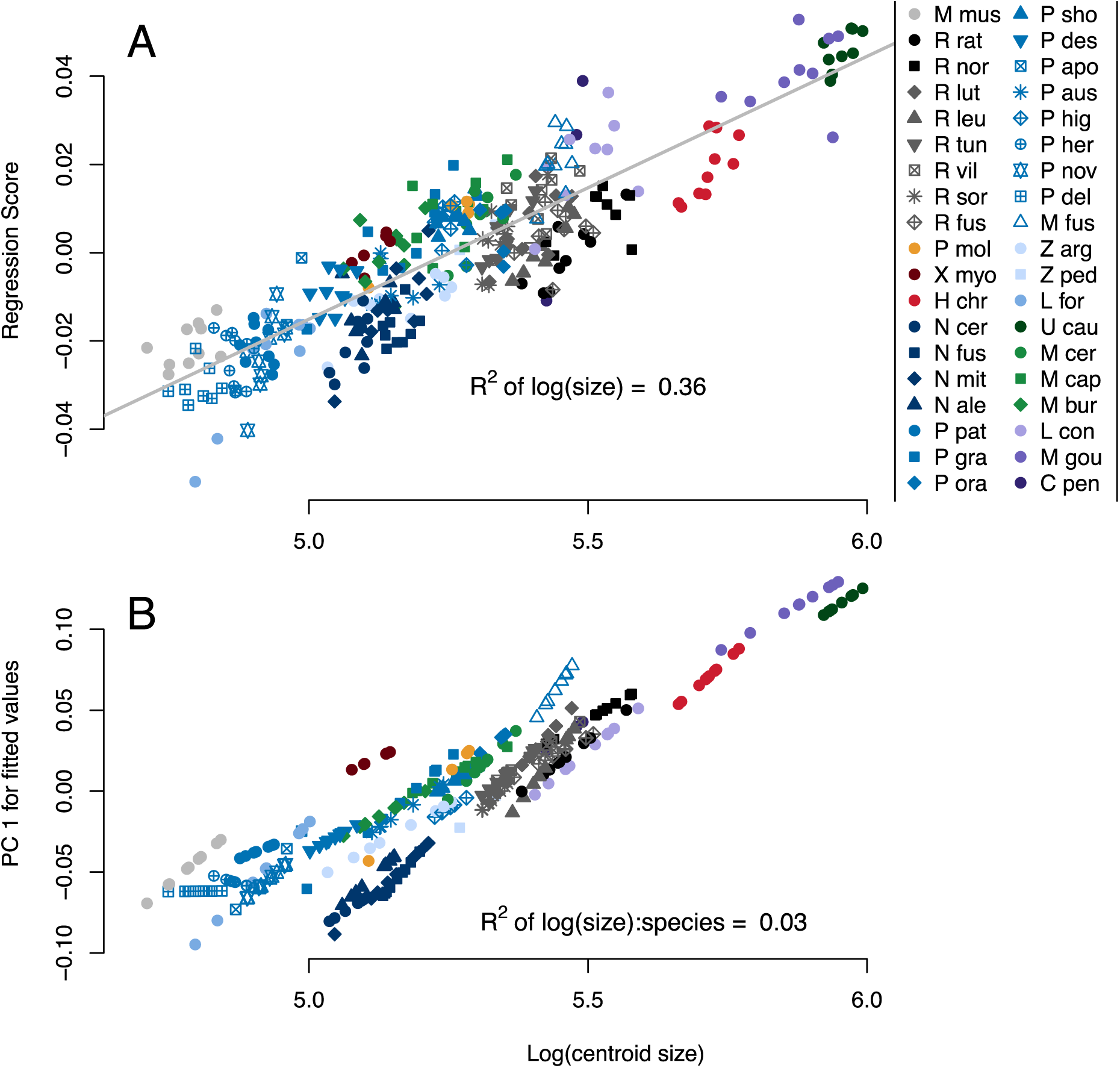
log centroid size, versus the regression scores of shape on size for each specimen (A). The overlaid grey line represents evolutionary allometry, or a regression on the mean specimen from every static allometry (Cheverud 1982) (see fig. 3). Predicted values for each species highlighting similarities in static allometric slopes (B).

Second, variation in evolutionary (among clades) allometries was tested using an ANCOVA model similar to the above (see table 1*B*). Howevever, instead of species, six major clades were defined as a radiation from an ancestor that arrived in Australia at a distinct time, after Aplin and Ford (2014) plus monospecific lineages for house mouse *Mus musculus* (Linneaus 1758) and prehensile-tailed mouse *P*. *mollipilosis* (see fig. 1). We also compared these results with a phylogenetic ANCOVA (pANCOVA) using *geomorph*’s *procD*.*pgls* (Adams 2014), which excutes the ANCOVA model in a phylogenetic framework (see table A5). This pANCOVA used mean centroid sizes from all 37 species included in the tree.

One analytical challenge – even with our comparatively large sample sizes – is that available specimens per species may be too small to confidently estimate species-level allometric slopes. Therefore, we developed a new function, *rarefy*.*stat* in *landvR* (v.0.4) (Guillerme and Weisbecker 2019) and modified the *prop*.*parts* function from *ape* (v.5.2) (Paradis and Schliep 2018) to estimate how well our calculations of species-level allometric slopes withstood downsampling relative to the larger clade-level allometric slopes. We used this phylogeny-based rarefaction to assess whether our sampling effort could support our interpretations.

To produce figure 3, we first measured the observed allometric slope for every clade in figure 1, from the entire dataset to each individual species. Then we removed all but five random specimens (our smallest species sample size) from each clade and re-measured this rareified allometric slope, repeated 100 times. The median slope change between the random sample and the slope across the whole clade was calculated from the 100 values created by subtracting the observed slope from each rarefied slope in each clade. We calculated the absolute median slope change in degrees for each clade using the trigonometric formula for the angle between two slopes. We considered the rarefied slope to be significantly different to the observed slope if their angle was higher than 4.5° (5% of 90° – the largest possible angle between the two slopes). We visualized the results using a boxplot showing the 95% and 50% confidence intervals of the delta slope values and a scatterplot of the delta slope angles in context with the 4.5° confidence line (see fig. 3). To ensure our results were not biased by close phylogenetic relationships, we randomly assigned species into groups of the same size as the clades and reran the analysis above. We repeated this analysis for 100 different sets of random groups, ignoring single-species clades. Results were visualized using a boxplot of the median delta slopes (see fig. A2).

**Figure 3:**
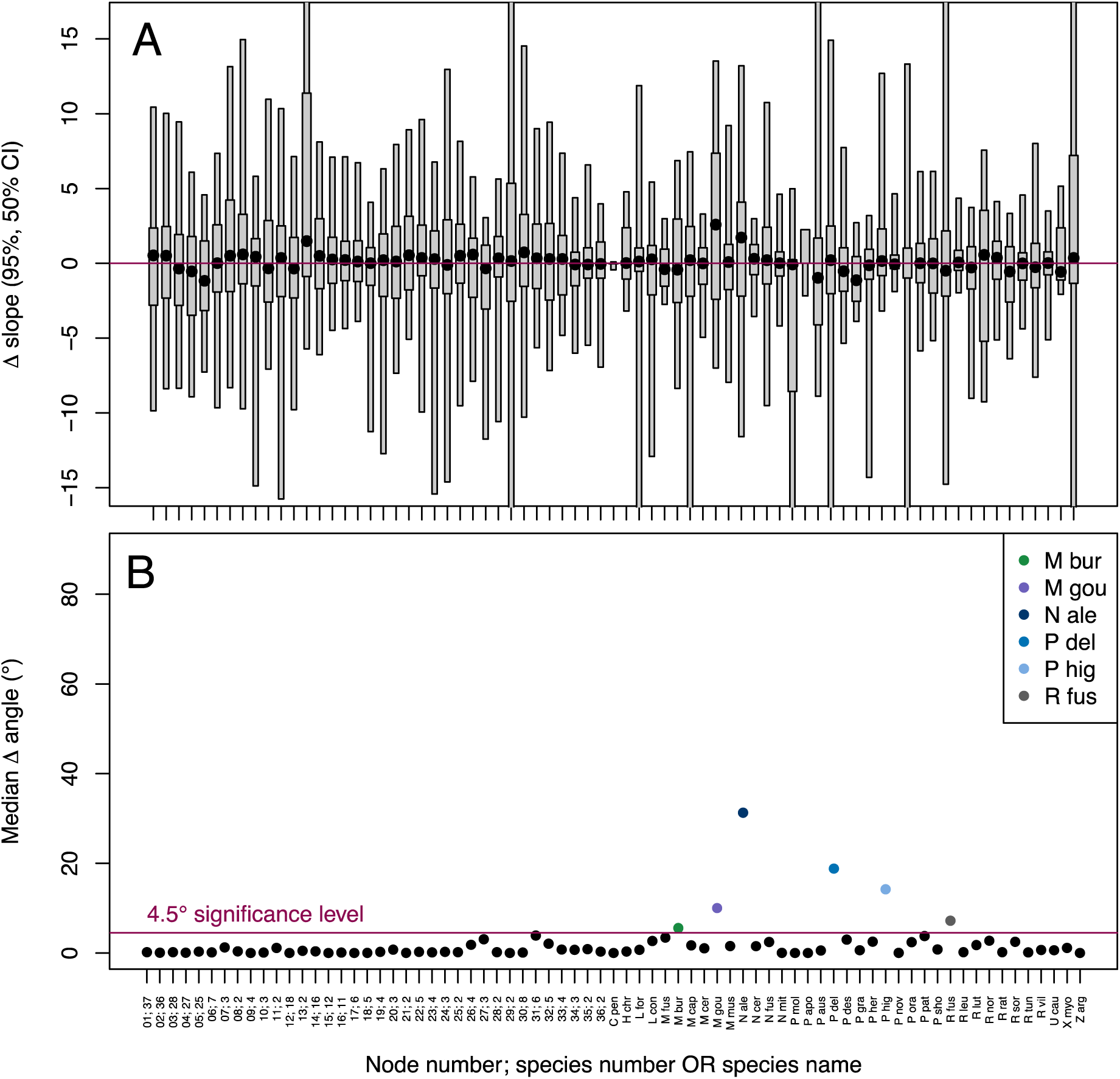
Phylogenetic rarefaction: boxplot showing confidence intervals (95%, 50%) and median value (black point) for each clade after 100 rarefaction replicates (A). Median delta slope angle (°) is the difference between the observed and rarefied slopes. Each point gives the clade’s median delta slope as compared to the 4.5° significant change in angle line (B). The x axis gives both the node number from figure 1 and the number of species included in that node from figure 1; monospecific clades have only species abbreviation. Outliers are colored and identified in the legend.

### CREA Shape Patterns

To address our second question on craniofacial evolutionary allometry (CREA) we assessed size and shape covariation using three types of plots. First, we used *geomorph*’s *procD*.*lm* to plot the evolutionary allometric relationship between log centroid size versus the regression of shape on size (see fig. 4*A*) (Drake and Klingenberg 2008). Second, we used *geomorph*’s *plotTangentSpace* to plot a principal components analysis (PCA) to provide a “size-less” morphospace comparison of the mean shapes for each species (fig. 4*B*). Third, we visualized the cranial shape variation across the minimum and maximum values of principal component (PC)1 using landmark heatmaps produced by *landvR* function *procrustes*.*var*.*plot* (fig. 4*C-F*) (Guillerme and Weisbecker 2019; Weisbecker et al. 2019). The heatmaps allowed us to determine whether the shape variation pattern resembled CREA (Cardini et al. 2015).

**Figure 4:**
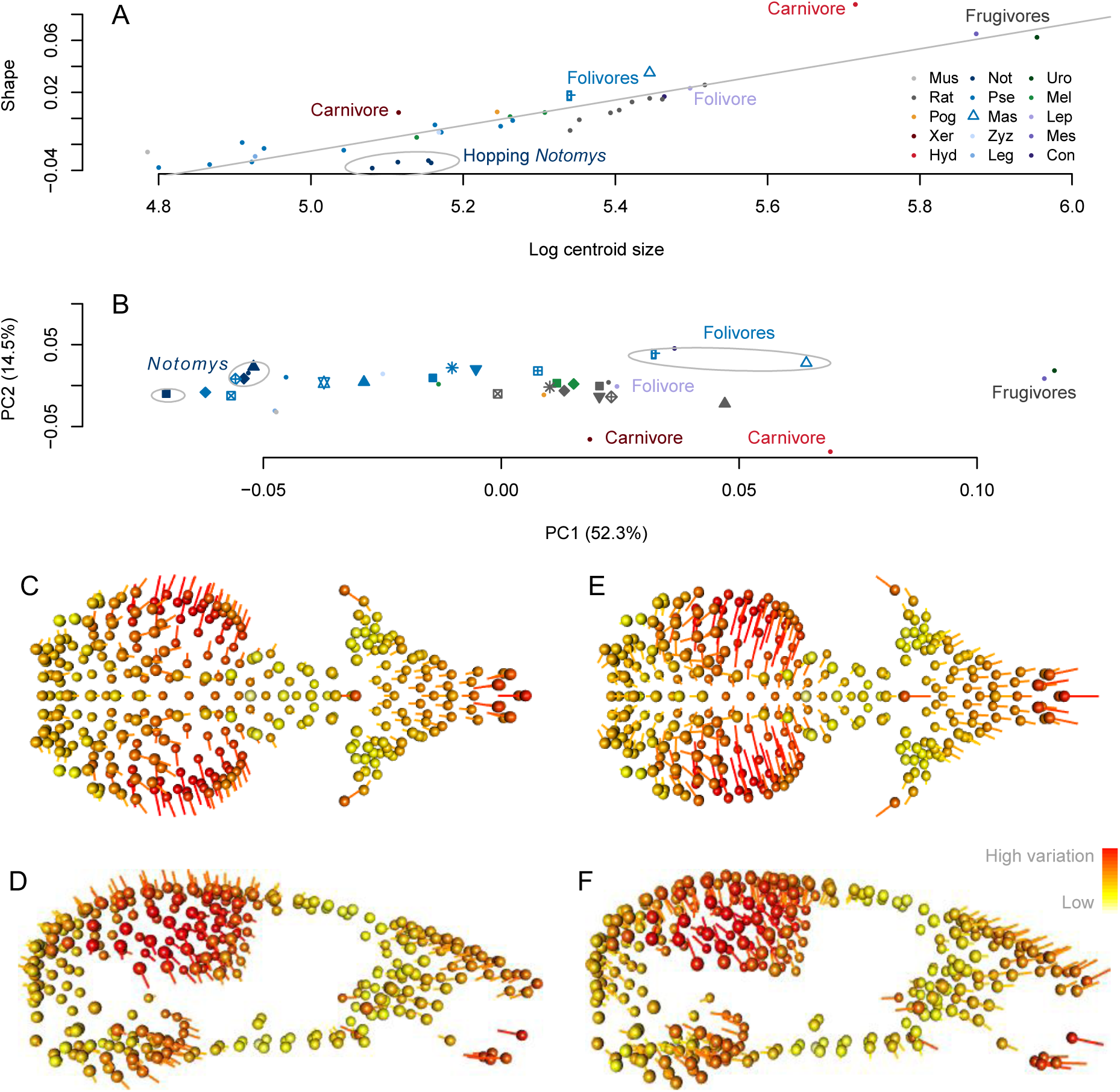
Evolutionary allometric variation: plot of log centroid size versus the projected regression score of allometric shape on size (A). The grey regression line indicates the common evolutionary allometry trajectory. Labels and “hand-drawn” ellipses indicate species sharing a diet or locomotion specialization. PCA plot of PC1 and PC2 separates mean shapes of many specialists from the main cluster (B). Legend as in figure 2. Landmark heatmaps of shape change from the mean shape to PC1 minimum, dorsal view (C) and lateral view (D). Landmark heatmaps from mean shape to the PC1 maximum, dorsal view (E) and lateral view (F). Spheres show landmark positions for the mean shape and vectors show the direction of change to the extreme PC1 shapes.

### Rates of Shape Evolution

To address our third question on stabilizing selection and facilitation, we used an evolutionary allometry plot (fig. 4*A*) to identify two types of outliers: large-bodied specialists on the common allometric line as well as specialists diverging from it. While specialists for frugivory, carnivory, and hopping locomotion are relatively easy to define, folivores exist along a spectrum. We identified our three “specialist folivore” species based on descriptions of cranial modifications for folivory and field studies demonstrating diet dependence: the broad-toothed rat, *Mastacomys fuscus*, the Hastings River mouse, *Pseudomys oralis* (Thomas, 1921), and the greater stick-nest rat, *Leporillus conditor* (Sturt, 1848) (Watts and Braithwaite 1978; Fox et al. 1994; Murray et al. 1999; Ryan, Moseby, and Paton 2003; Breed and Ford 2007). Using the mean shapes and the phylogeny of 37 species we ran *geomorph*’s *compare*.*evol*.*rates* to find pairwise comparisons of shape evolution rates between specialists and between specialists to non-specialists (see table A6). The Bonferroni correction accounted for multiple comparisons (Bonferroni et al. 1936).

All specimen surface files for Australian rodent crania included in this study are publicly available for unrestricted download from MorphoSource project 561 (http://www.morphosource.org/Detail/ProjectDetail/Show/project_id/561). All other data, including museum metadata, landmarking coordinates, and the molecular phylogeny, are publicly available on GitHub in the Data folder. Our GitHub repository also contains the R scripts needed to reproduce all analyses, tables, and figures (https://github.com/miracleray/allometry-rodents).

## Results

### 1) Allometric variation

At the static allometry level, the ANCOVA indicates that size accounts for a large fraction (36.5%) of shape (R_2_ = 0.365, p < 0.002), only slightly less than the variation explained by species affiliation (R_2_ = 0.405, p < 0.002) (table 1*A*). The post-hoc test for homogeneity of slopes found that, out of 703 pairwise comparisons, only nine had significant differences in slopes (table A4). The New Holland mouse, *Pseudomys novaehollandiae* (Waterhouse, 1843) had the greatest number of significant pairwise differences with six (out of a possible 37). All other species with significant pairwise differences had less than three such comparisons (table A4). Together, the ANCOVA and the homogeneity of slopes tests reject the extrinsic pressure hypothesis for ecological opportunity in Australia. Instead they support conserved allometry in which murid rodent species have parallel static allometric slopes (fig. 2).

Second, the evolutionary allometry (among clades) ANCOVA also showed a high R_2_ term for size (R_2_ = 0.364, p < 0.002), about twice that of clade (R_2_ = 0.175, p < 0.002), indicating a conserved allometric signal across the phylogeny (table 1*B*). The ANCOVA also revealed a small yet significant interaction term between clade and log centroid size (table 1*B*). This interaction term means that evolutionary allometric slopes differ slightly among clades. However, the coefficient of determination (R_2_) of this interaction is small: it only accounted for 5.6% of variation, compared with 37% and 18% for log centroid size and clade, respectively (table 1*B*). The pANCOVA of mean shapes against size returned similar results, with size accounting for 41% of variation. While the interaction term (table A5, R_2_ = 0.134, p < 0.02) is higher in this analysis, it uses fewer data points (mean shapes) within a phylogenetic context. Note that in both analyses, the species-rich Pseudomys division (n = 19 species) may introduce some sampling bias relative the other clades (n = 1-6 species).

Figure 2*A* illustrates the evolutionary allometry (grey line), which is shallower than static allometries but is correlated with the overall trend of size and shape across all species (table 1*A*, R_2_= 0.36, p < 0.002). This pattern occurs when slopes stay constant and species vary only slightly in y-intercepts (Pélabon et al. 2014). Here, y-intercepts generally decrease with increasing body size, which generates the shallower evolutionary allometry slope.

Third, phylogenetic rarefaction supports our that sampling is sufficient to reject a hypothesis of non-parallel slopes by showing that the conserved allometric trends found at the species and clade levels persist at a low sample size (n = 5) across each node of the tree (fig. 3). All clades had a median delta slope change less than 2.6 relative to the all-clade slope (fig. 3*A*), when converted to degrees, this corresponds to 93% of clades (67 of 73) remaining under the conservative 4.5° cut-off for slope angle change (fig. 3*B*). Randomizing the phylogeny did not change these results (fig. A2). Larger clades have much larger sample sizes to begin with, yet their median slope angles did not change significantly when downsampled. Therefore, we conclude that sample sizes of 5 or greater are sufficient for our study.

### 2) Craniofacial Evolutionary Allometry (CREA)

Consistent with the ANCOVAs, the evolutionary allometry plot shows few species diverging from the common evolutionary allometric trajectory (fig. 4*A*), establishing that a conserved pattern of cranial allometry exists in Australian rodents. The first two PC axes of the PCA represent 67% of the mean species shape variation (52.3% PC1, 14.5% PC2) while remaining PCs each explained 8.0% of variation or less (the first 10 PCs had a proportion of variance >1% each). Most of the shape variation, as identified by PC1 (fig. 4*B*), relates to allometry, with most species falling in the same order along the x-axes of centroid size and PC1 (fig. 4*A,B*). The PC1 landmark heatmaps clearly illustrate the PC1 minimum cranium having a larger basicrania and shorter snout compared to the mean shape (fig. *4C,D*) and the PC1 maximum cranium showing the opposite trend (fig. *4E,F*). These shapes are fully consistent with CREA (Cardini and Polly 2013; Cardini et al. 2015; Tamagnini et al. 2017).

Specialist species that diverge from the allometry plot also diverge from the main cluster of more generalist species along PC2 in the PCA (fig. 4*A,B*). Folivorous specialists score highest on PC2 (fig. 4*B*, dark purple circle, blue open triangle and quartered circle) while carnivorous specialists score lowest on PC2 (fig. 4*B*, dark red and red circles).

### 3) Rates of shape evolution on and off the common evolutionary allometric trajectory

Two frugivores – the black-footed tree rat, *Mesembriomys gouldi* (Palmer, 1906) and the giant white-tailed rat, *Uromys caudimaculatus* (Krefft, 1867) – independently evolved large bodies and outlying shapes along PC1, doing so on the common evolutionary allometric trajectory (fig. 4*A,B*). Of the three folivores, only the Hastings River mouse, *P*. *oralis* and the broad-toothed rat, *M*. *fuscus*, diverge along PC2 and from the common evolutionary allometric trajectory (fig. 4). The third folivore, the greater stick-nest rat, *L*. *conditor* falls directly along the allometric trajectory (fig. 4). Both carnivores diverge along common evolutionary allometry trajectory and along PC2 with the opposite loading from the folivores (fig. 4*A,B*). The water rat, *Hydromys chrysogaster* (Geoffrey, 1804) appears most divergent from the common evolutionary allometry trajectory (fig. 4*A*). The bipedal hopping *Notomys* appear to have an among-clade allometry that diverged in y-intercept but not in slope from other, predominantly quadrupedal Australian rodents (fig. 4*A*). They consistently show low PC1 scores (fig. 4*B*).

Pairwise analysis of shape evolution rates revealed that crania of large-bodied frugivores evolved 3.96 times faster than those of non-specialists (table A6, p = 0.04). The two frugivores evolved on the common evolutionary allometric trajectory independently, supporting the hypothesis for facilitation along a line of least resistance, an outcome of stabilizing selection. All other pairwise comparisons were non-significant, including for specialists diverging from the common evolutionary allometry trajectory (table A6, p > 0.05).

## Discussion

We find strong, conserved allometry of skull shape across nearly all levels of the Australian murid rodent phylogeny, explaining substantial amounts of the variation (roughly 40% of both the static (species-level) and evolutionary variation as well as over half (52%) of variation along PC1). We therefore find no support for the extrinsic pressure hypothesis (that there should be divergence of allometric slopes because of divergent selection pressures). In fact, with very few exceptions, all species retain a similar allometric slope across divergences as wide as ten million years – since the split between *Mus* and the clade including all native Australian rodents (Aghová et al. 2018). Our new phylogeny-based rarefaction, bootstrapping, and randomization method shows that this allometric conservation transcends taxanomic boundaries across the entire sample, with nearly no significant differences between static and evolutionary allometric slopes. Indeed, static allometric slope angles showed almost no significant changes between samples, even when species from different clades were combined at random. The strict conservation of allometric scaling is particuarly striking for such a speciose group encompassing six major radiations onto a new continent with novel environments (Yoder et al. 2010; Aplin and Ford 2014). While strongly conserved allometry has been detected among closely related species (Singleton 2002; Cardini et al. 2015; Munds et al. 2018), we are not aware of similar levels of allometric conservation across any other large radiation of mammals. Our results therefore demonstrate rodents to be an example of extreme allometric conservatism within the placentals, a clade otherwise thought to have a high degree of evolvability in cranial allometry (Tsuboi et al. 2018).

Our heatmap visualizations of both allometric and ordinated (PCA) shape variation demonstrate that the high degree of allometry in Australian murids coincides with shape variation known as “craniofacial evolutionary allometry” (CREA). CREA is found across diverse mammalian lineages, and describes allometric shape variation where larger species have relatively longer snouts and smaller braincases compared to related species with smaller body sizes (Cardini and Polly 2013; Cardini 2019). However, due to their particularly conserved allometry, Australian murid rodents appear to be uniquely constrained to CREA compared to other mammals.

The underlying cause of CREA across Mammalia is still under investigation (Cardini 2019). Current hypotheses include developmental constraints as well as persistent selection on function via stabilizing selection (Cardini and Polly 2013). The instrinsic constraint hypothesis is certainly supported by the finding that murid rodents, with fast reproduction and altricial neonates compared to other placentals, would have shape evolution driven primarily by size (Porto et al. 2013). Furthermore, Australian murids vary in reproductive rate by clade, with the highest reproductive rates occuring in the most morphologically conserved clade of native *Rattus* (Yom-Tov 1985; Geffen et al. 2011; Rowe et al. 2011). Therefore, our results position Australian murid rodents as potentially developmentally-constrained exceptions to the otherwise developmentally-diverse placentals, supporting the general hypothesis for increased morphological diversity in placentals that have longer relative gestation times than rodents (Porto et al. 2013; Tsuboi et al. 2018).

Despite the strong indication of a developmental constraint, constraint hypotheses are not mutally exclusive with hyptheses of stabilizing selection and we found complimentary lines of evidence that support a strong role of stabilizing selection.In particular, stabilizing selection can act on available genetic variation to produce an allometric line of least resistance that scales viable and functional morphological ratios with body size (Schluter 1996). In our dataset, two frugivores from different radiations evolved large body sizes with similar cranial shapes that sit along the evolutionary allometry trajectory; this was accompanied by significantly faster rates of shape evolution compared to non-specialists. Faster evolution is predicted under the stabilizing selection hypothesis because of facilitation by the allometric line of least resistance (Schluter 1996; Marroig and Cheverud 2005). This appears to be a likely scenario for Australian large-bodied frugivores because experimental work has suggested that the general rodent gnawing apparatus maintains frugivory with few or no changes (Cox et al. 2012; Maestri et al. 2016).

Folivores and carnivores may also make a case for the existence of stabilizing selection in Australian murids. These two groups deviated from the common allometric line and in the PCA. Folivores showed higher PC2 values corresponding to broader molars than non-folivorous species of the same size. Carnivores showed lower PC2 scores, with fewer teeth and a rostrum morphology adaptive for capturing prey; an unusual niche for rodents (Fabre et al. 2017). These morphological changes did not alter the conserved species-level allometric slope, even for carnivorous water rat, *Hydromys chrysogaster*, whose mean projected shape to size ratio falls noticeably above the common evolutionary allometric trajectory. It is possible that adaptations away from the common evolutionary allometric line come with trade offs. For example, a previous anatomical study of *H*. *chrysogaster* (Fabre et al. 2017) suggested that they maximize bite force by reducing movement at their craniomandibular joint, but that this causes maladaptive patterns of tooth microwear and broken incisor tips (Druzinsky 2015; Fabre et al. 2017). This trade-off suggests that disruptive selection occurs on Australian murids, but that stabilizing selection acts as a strong antagonist to changes away from the evolutionary allometric line.

Australian murid rodents can be compared to many other mammalian radiations with regards to allometry and conserved morphology. For example, Indo-Australian murid rodents evolved carnivory five times (Rowe et al. 2016) and south-east Asian murid vermivores evolved unusual crania that appear to diverge from CREA (Esselstyn et al. 2012; Rickart et al. 2019). These relatives could be used to explore how disruptive or directional selection could overpower previously existing stabilizing selection. Indeed, the intense stabilizing selection that we infer acts on the complex gnawing apparatus of rodents invites and comparisons to the unrelated clade of multituberculates, which share features of this apparatus (Lazzari et al. 2010). This combination of characters appears to correspond with similar patterns of low cranial diversity, high species richness, and success in a range of environments (Lazzari et al. 2010). Indeed, clades with low morphological diversity could have a highly adaptive suite of morphological ratios whose biomechanics scale along an allometric line (Marroig and Cheverud 2005; Cardini and Polly 2013). In these cases, non-allometric morphological diversity would be determined by intrinsic constraints and how much deviation from existing allometry is tolerated by stabilizing selection (Estes and Arnold 2007). New World monkeys show allometric patterns suggestive of both constraint and stabilizing selection, with evidence that the latter could have facilitated evolution along a line of least resistance (Marroig and Cheverud 2005, 2010). This clade would provide an ideal comparison to altricial Australian murids to disentangle these two factors further because monkeys – unlike murids – have slow reproductive rates, like most other placental clades (Lillegraven 1974; Porto et al. 2013; Tsuboi et al. 2018).

## Conclusions

Understanding the specific roles of intrinsic constraints and stabilizing selection on conserved allometric patterns like CREA has the potential to answer fundamental macroevolutionary questions (Cardini 2019). However, the conceptual difference is difficult to disentangle because, as our study shows, CREA appears to be a long-term emergent property of both genetics and selection (i.e. it represents the compromise between instrinsic developmental programs and extrinsic selection on viable forms throughout ontogeny) (Pélabon et al. 2014; Brigandt 2015). Measuring ontogenetic allometry could eliminate intrinsic constraints as the limiting factor if high ontogenetic variation exists, indicating a larger role for stabilizing selection (Jamniczky and Hallgrímsson 2009). There is already some evidence that murid rodents have highly variable ontogenetic allometry despite conserved static allometries (Wilson and Sánchez-Villagra 2009). Finally, the trade-off observed between conserved allometric shape and orthogonal shape variation deserves further exploration as a possible avenue to understand the interaction between factors influencing allometry and total morphological variation.

## Abbreviations

CREA: Craniofacial Evolutionary Allometry
PCA: Principal component analysis
PC: Principal component
3D: Three dimensional

## Acknowledgements

We thank Dr Heather Janetzki for hosting AEM many times in the mammal collections at the Queensland Museum, Laura Cook for hosting at the Museum Victoria, Dr Sandy Ingleby for hosting at the Australian Museum, and Dr David Stemmer for loaning specimens from the South Australian Museum. Thanks to lab assistants Lauren Thornton and Aubrey Keirnan for help uploading 3D meshes to Morphosource. We also thank Editor Dan Rabosky and two anonymous reviewers for their insightful comments that greatly enhanced the manuscript. This work was funded by Discovery Grant DP170103227 to VW and MJP, as well as by a School of Biology Postgraduate Travel grant to AEM.

