## Supplementary material for "Australian rodents reveal conserved Cranial Evolutionary Allometry across 10 million years of murid evolution": table A1

**Table A1:** sex and specimen numbers by species. Museum abbreviations are as follows: Australian Museum (AM), Museums Victoria (MV), Queensland Museum (QM), and South Australian Museum (SAM). Catalogue numbers match those exactly in respective museum databases. Genus and species are given in three letter abbreviations, which match those used throughout the article.

| **Museum** | **CatNum** | **Genus** | **Species** | **Sex** |
| --- | --- | --- | --- | --- |
| AM | M27465 | Lep | con | Female |
| AM | S1600 | Lep | con | Female |
| AM | M27463 | Lep | con | Male |
| AM | M3061 | Lep | con | Male |
| AM | M3063 | Lep | con | Male |
| AM | M3065 | Lep | con | Male |
| AM | S1597 | Lep | con | Male |
| AM | M3409 | Lep | con | Unknown |
| AM | M3457 | Lep | con | Unknown |
| AM | M10765 | Mas | fus | Female |
| AM | M7170 | Mas | fus | Female |
| AM | M36984 | Mas | fus | Male |
| AM | M27895 | Mes | gou | Male |
| AM | M8345 | Mes | gou | Male |
| AM | M27887 | Not | ale | Female |
| AM | M5688 | Not | ale | Female |
| AM | M10005 | Not | ale | Male |
| AM | M23514 | Not | cer | Female |
| AM | M8496 | Not | cer | Female |
| AM | M23515 | Not | cer | Male |
| AM | M23516 | Not | cer | Male |
| AM | M8493 | Not | cer | Male |
| AM | M8495 | Not | cer | Male |
| AM | M8498 | Not | cer | Male |
| AM | M23644 | Not | fus | Female |
| AM | M23646 | Not | fus | Female |
| AM | M23508 | Not | fus | Male |
| AM | M4860 | Not | fus | Male |
| AM | M4858 | Not | mit | Female |
| AM | M4907 | Not | mit | Female |
| AM | M5028 | Not | mit | Female |
| AM | M8617 | Not | mit | Female |
| AM | M3735 | Not | mit | Male |
| AM | M4853 | Not | mit | Male |
| AM | M8192 | Not | mit | Male |
| AM | M10007 | Pse | aus | Female |
| AM | M10008 | Pse | aus | Female |
| AM | M10009 | Pse | aus | Male |
| AM | M44869 | Pse | aus | Male |
| AM | M9673 | Pse | gra | Female |
| AM | M31106 | Pse | gra | Male |
| AM | M9945 | Pse | gra | Male |
| AM | M26079 | Pse | her | Female |
| AM | M9207 | Pse | her | Female |
| AM | M25376 | Pse | her | Male |
| AM | M25377 | Pse | her | Male |
| AM | M29421 | Pse | her | Male |
| AM | M32487 | Pse | her | Male |
| AM | M4872 | Pse | hig | Female |
| AM | M4876 | Pse | hig | Female |
| AM | M2468 | Pse | hig | Female |
| AM | M4874 | Pse | hig | Male |
| AM | M4875 | Pse | hig | Male |
| AM | M4877 | Pse | hig | Male |
| AM | M4878 | Pse | hig | Male |
| AM | M12553 | Pse | nov | Female |
| AM | M12557 | Pse | nov | Female |
| AM | M25630 | Pse | nov | Female |
| AM | M9155 | Pse | nov | Female |
| AM | M12544 | Pse | nov | Male |
| AM | M8938 | Pse | nov | Male |
| AM | M9150 | Pse | nov | Male |
| AM | M25633 | Pse | ora | Female |
| AM | M29840 | Pse | ora | Female |
| AM | M12346 | Pse | ora | Male |
| AM | M6529 | Xer | myo | Female |
| AM | M9991 | Xer | myo | Female |
| AM | M10434 | Xer | myo | Male |
| AM | M6868 | Xer | myo | Male |
| AM | M21205 | Zyz | arg | Female |
| AM | M21225 | Zyz | arg | Male |
| AM | M24766 | Zyz | arg | Male |
| MV | C31802 | Mas | fus | Female |
| MV | C31804 | Mas | fus | Female |
| MV | C31805 | Mas | fus | Female |
| MV | C15025 | Mas | fus | Male |
| MV | C15027 | Mas | fus | Male |
| MV | C15034 | Mas | fus | Male |
| MV | C25854 | Mas | fus | Male |
| MV | DTC465 | Mel | cap | Female |
| MV | DTC462 | Mel | cap | Male |
| MV | DTC464 | Mel | cap | Male |
| MV | DTC466 | Mel | cap | Male |
| MV | C7570 | Mes | gou | Female |
| MV | C7571 | Mes | gou | Male |
| MV | C36854 | Not | ale | Female |
| MV | C36848 | Not | ale | Male |
| MV | C36885 | Not | ale | Male |
| MV | C36886 | Not | ale | Male |
| MV | C36892 | Not | ale | Male |
| MV | C7356 | Not | cer | Female |
| MV | C9081 | Not | cer | Female |
| MV | C7833 | Not | fus | Female |
| MV | C7834 | Not | fus | Female |
| MV | C7835 | Not | fus | Male |
| MV | C7836 | Not | fus | Male |
| MV | C15045 | Not | mit | Female |
| MV | C157 | Pse | aus | Female |
| MV | C4886 | Pse | aus | Male |
| MV | C4883 | Pse | aus | Unknown |
| MV | C4884 | Pse | aus | Unknown |
| MV | C10770 | Pse | des | Female |
| MV | C36860 | Pse | des | Female |
| MV | C36878 | Pse | des | Female |
| MV | C10755 | Pse | des | Male |
| MV | C36870 | Pse | des | Male |
| MV | C36879 | Pse | des | Male |
| MV | C18705 | Pse | sho | Female |
| MV | C19922 | Pse | sho | Female |
| MV | C22118 | Pse | sho | Female |
| MV | C19925 | Pse | sho | Male |
| MV | C19926 | Pse | sho | Male |
| MV | C22111 | Pse | sho | Male |
| MV | C37144 | Pse | sho | Male |
| MV | C37172 | Pse | sho | Male |
| MV | C20261 | Rat | nor | Female |
| MV | C7338 | Rat | nor | Female |
| MV | C29709 | Rat | nor | Male |
| MV | C33155 | Rat | nor | Male |
| MV | C6982 | Uro | cau | Female |
| MV | C6983 | Uro | cau | Female |
| MV | C6984 | Uro | cau | Male |
| QM | J10055 | Hyd | chr | Female |
| QM | J16824 | Hyd | chr | Female |
| QM | J9835 | Hyd | chr | Female |
| QM | J9944 | Hyd | chr | Female |
| QM | J21273 | Hyd | chr | Female |
| QM | J10127 | Hyd | chr | Male |
| QM | J11137 | Hyd | chr | Male |
| QM | J15330 | Hyd | chr | Male |
| QM | J17593 | Hyd | chr | Male |
| QM | J7889 | Hyd | chr | Male |
| QM | JM11585 | Leg | for | Female |
| QM | JM19648 | Leg | for | Female |
| QM | JM3736 | Leg | for | Female |
| QM | J14755 | Leg | for | Male |
| QM | JM12328 | Leg | for | Male |
| QM | JM10913 | Leg | for | Unknown |
| QM | JM14441 | Leg | for | Unknown |
| QM | JM14820 | Leg | for | Unknown |
| QM | JM4955 | Leg | for | Unknown |
| QM | J19997 | Mel | bur | Female |
| QM | J4808 | Mel | bur | Female |
| QM | J9801 | Mel | bur | Female |
| QM | JM13946 | Mel | bur | Female |
| QM | JM15015 | Mel | bur | Female |
| QM | J10822 | Mel | bur | Male |
| QM | J17693 | Mel | bur | Male |
| QM | J9476 | Mel | bur | Male |
| QM | JM1301 | Mel | bur | Male |
| QM | JM3839 | Mel | bur | Male |
| QM | JM13959 | Mel | cap | Female |
| QM | JM13964 | Mel | cap | Unknown |
| QM | JM4224 | Mel | cap | Unknown |
| QM | JM4231 | Mel | cap | Unknown |
| QM | JM4233 | Mel | cap | Unknown |
| QM | JM4236 | Mel | cap | Unknown |
| QM | J22537 | Mel | cer | Female |
| QM | J3676 | Mel | cer | Female |
| QM | J8956 | Mel | cer | Female |
| QM | JM10991 | Mel | cer | Female |
| QM | J22096 | Mel | cer | Male |
| QM | J3675 | Mel | cer | Male |
| QM | J6502 | Mel | cer | Male |
| QM | J8960 | Mel | cer | Male |
| QM | JM6185 | Mel | cer | Male |
| QM | J19719 | Mes | gou | Female |
| QM | J16976 | Mes | gou | Male |
| QM | J19720 | Mes | gou | Male |
| QM | JM20901 | Mes | gou | Unknown |
| QM | J2562 | Mes | gou | Unknown |
| QM | J3078 | Mus | mus | Female |
| QM | J2993 | Mus | mus | Female |
| QM | J16129 | Mus | mus | Male |
| QM | J9161 | Mus | mus | Male |
| QM | JM1272 | Mus | mus | Male |
| QM | J13566 | Mus | mus | Male |
| QM | J16141 | Mus | mus | Male |
| QM | J16150 | Mus | mus | Male |
| QM | J3105 | Mus | mus | Unknown |
| QM | JM1027 | Mus | mus | Unknown |
| QM | J14754 | Not | ale | Male |
| QM | J10775 | Not | fus | Male |
| QM | J3351 | Not | mit | Male |
| QM | JM14683 | Pog | mol | Female |
| QM | JM10071 | Pog | mol | Male |
| QM | JM10590 | Pog | mol | Male |
| QM | JM8501 | Pog | mol | Male |
| QM | JM8841 | Pog | mol | Male |
| QM | JM11502 | Pse | del | Female |
| QM | JM12690 | Pse | del | Female |
| QM | JM12695 | Pse | del | Female |
| QM | JM12708 | Pse | del | Female |
| QM | JM18715 | Pse | del | Female |
| QM | JM11350 | Pse | del | Male |
| QM | JM19635 | Pse | del | Male |
| QM | JM2133 | Pse | del | Male |
| QM | JM8828 | Pse | des | Female |
| QM | JM14592 | Pse | des | Male |
| QM | JM4953 | Pse | des | Male |
| QM | JM15791 | Pse | gra | Female |
| QM | JM11213 | Pse | gra | Male |
| QM | JM14420 | Pse | gra | Male |
| QM | JM11182 | Pse | gra | Male |
| QM | JM19872 | Pse | gra | Unknown |
| QM | JM14849 | Pse | gra | Unknown |
| QM | JM1289 | Pse | her | Female |
| QM | JM2509 | Pse | her | Female |
| QM | JM5019 | Pse | her | Female |
| QM | J16683 | Pse | her | Male |
| QM | JM14455 | Pse | nov | Female |
| QM | J17920 | Pse | nov | Male |
| QM | JM14454 | Pse | nov | Male |
| QM | J20264 | Pse | ora | Female |
| QM | JM13548 | Pse | ora | Unknown |
| QM | JM13549 | Pse | ora | Unknown |
| QM | JM11008 | Pse | pat | Female |
| QM | JM11940 | Pse | pat | Female |
| QM | JM8654 | Pse | pat | Female |
| QM | JM10864 | Pse | pat | Male |
| QM | JM10865 | Pse | pat | Male |
| QM | JM12674 | Pse | pat | Male |
| QM | JM8830 | Pse | pat | Male |
| QM | JM12363 | Pse | pat | Male |
| QM | JM15010 | Pse | pat | Unknown |
| QM | J3488 | Pse | sho | Male |
| QM | J11226 | Rat | fus | Female |
| QM | J12672 | Rat | fus | Female |
| QM | J19105 | Rat | fus | Female |
| QM | JM12469 | Rat | fus | Female |
| QM | J10939 | Rat | fus | Male |
| QM | J3681 | Rat | fus | Male |
| QM | J9687 | Rat | fus | Male |
| QM | JM11916 | Rat | fus | Male |
| QM | JM15739 | Rat | fus | Male |
| QM | J10136 | Rat | leu | Female |
| QM | J8280 | Rat | leu | Female |
| QM | JM172014 | Rat | leu | Female |
| QM | JM2127 | Rat | leu | Female |
| QM | J10139 | Rat | leu | Male |
| QM | J10197 | Rat | leu | Male |
| QM | JM11803 | Rat | leu | Male |
| QM | JM17301 | Rat | leu | Male |
| QM | JM1768 | Rat | leu | Male |
| QM | J16918 | Rat | lut | Female |
| QM | J22555 | Rat | lut | Female |
| QM | J22595 | Rat | lut | Female |
| QM | J8922 | Rat | lut | Female |
| QM | J20340 | Rat | lut | Male |
| QM | J22819 | Rat | lut | Male |
| QM | J22885 | Rat | lut | Male |
| QM | JM14757 | Rat | lut | Male |
| QM | J22598 | Rat | lut | Male |
| QM | J11439 | Rat | nor | Female |
| QM | J17925 | Rat | nor | Female |
| QM | J10052 | Rat | nor | Male |
| QM | J17540 | Rat | nor | Male |
| QM | J17927 | Rat | nor | Male |
| QM | J10961 | Rat | rat | Female |
| QM | J20163 | Rat | rat | Female |
| QM | J3326 | Rat | rat | Female |
| QM | J4085 | Rat | rat | Female |
| QM | J16172 | Rat | rat | Male |
| QM | J17793 | Rat | rat | Male |
| QM | J17798 | Rat | rat | Male |
| QM | J17923 | Rat | rat | Male |
| QM | J6275 | Rat | rat | Male |
| QM | J17959 | Rat | sor | Female |
| QM | J22871 | Rat | sor | Female |
| QM | J3836 | Rat | sor | Female |
| QM | J8929 | Rat | sor | Female |
| QM | J17955 | Rat | sor | Male |
| QM | J20409 | Rat | sor | Male |
| QM | J9172 | Rat | sor | Male |
| QM | JM1313 | Rat | sor | Male |
| QM | JM9078 | Rat | sor | Male |
| QM | J16895 | Rat | tun | Female |
| QM | J22604 | Rat | tun | Female |
| QM | J9206 | Rat | tun | Female |
| QM | J9786 | Rat | tun | Female |
| QM | J22095 | Rat | tun | Male |
| QM | J22099 | Rat | tun | Male |
| QM | J9566 | Rat | tun | Male |
| QM | J9571 | Rat | tun | Male |
| QM | JM12504 | Rat | tun | Male |
| QM | J16963 | Rat | vil | Female |
| QM | J16964 | Rat | vil | Female |
| QM | J20160 | Rat | vil | Female |
| QM | J22613 | Rat | vil | Female |
| QM | J9162 | Rat | vil | Female |
| QM | J16967 | Rat | vil | Male |
| QM | J19057 | Rat | vil | Male |
| QM | J6719 | Rat | vil | Male |
| QM | J6721 | Rat | vil | Male |
| QM | J9682 | Rat | vil | Male |
| QM | J22538 | Uro | cau | Female |
| QM | JM2344 | Uro | cau | Female |
| QM | J22607 | Uro | cau | Female |
| QM | J11512 | Uro | cau | Male |
| QM | J20347 | Uro | cau | Male |
| QM | J9304 | Uro | cau | Male |
| QM | JM4924 | Xer | myo | Female |
| QM | JM2708 | Xer | myo | Male |
| QM | J22399 | Zyz | arg | Female |
| QM | JM12723 | Zyz | arg | Female |
| QM | J22398 | Zyz | arg | Male |
| QM | J22401 | Zyz | arg | Male |
| QM | JM14576 | Zyz | arg | Unknown |
| QM | JM14578 | Zyz | arg | Unknown |
| SAM | M1796 | Con | pen | Male |
| SAM | M4071 | Con | pen | Male |
| SAM | M392 | Con | pen | Unknown |
| SAM | M11646 | Pse | apo | Female |
| SAM | M13666 | Pse | apo | Male |
| SAM | M3468 | Pse | apo | Unknown |
| SAM | M4379 | Zyz | ped | Female |
| SAM | M2412 | Zyz | ped | Unknown |
