## Supplementary material for "Australian rodents reveal conserved Cranial Evolutionary Allometry across 10 million years of murid evolution": table A2

**Table A2:** landmark definitions and point counts. Names indicate type of landmark and location (see figure A1). LM refers to a fixed landmark, Curve refers to the semi-landmarks placed on the same user-defined curve, and Patch refers to the patch semi-landmarks placed on the same surface and bounded by the same set of semi-landmark curves.

| **Name** | **Definition** | **Points included** |
| --- | --- | --- |
| LM1 | Anterior most point of nasal along central suture | 1 |
| LM2 | Central intersection of nasal and frontal bones | 1 |
| LM3 | Central intersection of frontal and parietal | 1 |
| LM4 | Central intersection of parietal and interpaterietal | 1 |
| LM5 | Central intersection interpaterietal and occipital | 1 |
| LM6 | Dorsal and central most point of the foramen magnum | 1 |
| LM7 | Ventral and central most point of the foramen magnum | 1 |
| LM8 | Central intersection of the basioccipital and basisphenoid suture | 1 |
| LM9 | Central and posterior most point of palatine | 1 |
| LM10 | Centerpoint between the posterior most tips of the anterior palantine foramen | 1 |
| LM11 | Centerpoint between the anterior most tips of the anterior palantine foramen | 1 |
| LM12 | Posterior point of incisor alveolar margin with center of the incisor (right) | 1 |
| LM13 | Posterior point of incisor alveolar margin with center of the incisor (left) | 1 |
| LM14 | Anterior and ventral most point of intersection between the premaxilla and incisor (right) | 1 |
| LM15 | Anterior and ventral most point of intersection between the premaxilla and incisor (left) | 1 |
| LM16 | Anterior and lateral most point of the premaxillar / nasal suture (right) | 1 |
| LM17 | Anterior and lateral most point of the premaxillar / nasal suture (left) | 1 |
| LM18 | Anterior most point of the infraorbital foramen (right) | 1 |
| LM19 | Anterior most point of the infraorbital foramen (left) | 1 |
| LM20 | Lateral edge of infraorbital foramen and anterior tip of zygomatic arch (right) | 1 |
| LM21 | Lateral edge of infraorbital foramen and anterior tip of zygomatic arch (left) | 1 |
| LM22 | Most posterior point of supraorbital (right) | 1 |
| LM23 | Most posterior point of supraorbital (left) | 1 |
| LM24 | Posterior intersection of the premaxilla and maxilla (right) | 1 |
| LM25 | Posterior intersection of the premaxilla and maxilla (left) | 1 |
| LM26 | Intersection of the frontal, squamosal, and parietal bones (right) | 1 |
| LM27 | Intersection of the frontal, squamosal, and parietal bones (left) | 1 |
| LM28 | Anterior most point of intersection of posterior zygomatic arch with squamosal (right) | 1 |
| LM29 | Anterior most point of intersection of posterior zygomatic arch with squamosal (left) | 1 |
| LM30 | Posterior most point of intersection of posterior zygomatic arch with squamosal (right) | 1 |
| LM31 | Posterior most point of intersection of posterior zygomatic arch with squamosal (left) | 1 |
| LM32 | Intersection of parietal, squamosal, and occipital sutures (right) | 1 |
| LM33 | Intersection of parietal, squamosal, and occipital sutures (right) | 1 |
| LM34 | Intersection of parietal and occipital bones (right) | 1 |
| LM35 | Intersection of parietal and occipital bones (left) | 1 |
| LM36 | Lateral occipital condyle intersect with the edge of the foramen magnum (right) | 1 |
| LM37 | Lateral occipital condyle intersect with the edge of the foramen magnum (left) | 1 |
| LM38 | Paraoccipital process (right) | 1 |
| LM39 | Paraoccipital process (left) | 1 |
| LM40 | Lingual tip of bulla (right) | 1 |
| LM41 | Lingual tip of bulla (left) | 1 |
| LM42 | Posterior point of pterygoid process (right) | 1 |
| LM43 | Posterior point of pterygoid process (left) | 1 |
| LM44 | Posterior most point of tooth row (right) | 1 |
| LM45 | Posterior most point of tooth row (left) | 1 |
| LM46 | Anterior most point of tooth row (right) | 1 |
| LM47 | Anterior most point of tooth row (left) | 1 |
| LM48 | Posterior most point of maxilla contribution to zygomatic arch (right) | 1 |
| LM49 | Posterior most point of maxilla contribution to zygomatic arch (left) | 1 |
| LM50 | Posterior point of external auditory meatus (right) | 1 |
| LM51 | Posterior point of external auditory meatus (left) | 1 |
| LM52 | Dorsal most point of external auditory meatus (right) | 1 |
| LM53 | Dorsal most point of external auditory meatus (left) | 1 |
| LM54 | Anterior and ventral most point of external auditory meatus (right) | 1 |
| LM55 | Anterior and ventral most point of external auditory meatus (left) | 1 |
| LM56 | Posterior and dorsal most point of intersection between the dentary and squamosal (right) | 1 |
| LM57 | Posterior and dorsal most point of intersection between the dentary and squamosal (left) | 1 |
| LM58 | Anterior and lateral most point of the zygomatric arch (right) | 1 |
| LM59 | Anterior and lateral most point of the zygomatric arch (left) | 1 |
| LM60 | Central anterior ventral most point of nasal opening | 1 |
| Curve1 | Central nasal suture between LM1 and 2 | 3 |
| Curve2 | Central frontal suture between LM2 and 3 | 4 |
| Curve3 | Central parietal suture between LM3 and 4 | 2 |
| Curve4 | Central interparietal line between LM4 and 5 | 1 |
| Curve5 | Central occipital line between LM5 and 6 | 2 |
| Curve6 | Central basisphenoid suture between LM7 and 8 | 2 |
| Curve7 | Central line of hard palate between LM9 and 10 | 3 |
| Curve8 | Nasal and premaxilla suture between LM16 and 2 (right) | 5 |
| Curve9 | Nasal and premaxilla suture between LM15 and 2 (left) | 5 |
| Curve10 | Lateral most edge of frontal between LM2 and 26 (right) | 3 |
| Curve11 | Lateral most edge of frontal between LM2 and 27 (left) | 3 |
| Curve12 | Frontal and parietal suture between LM3 and 26 (right) | 3 |
| Curve13 | Frontal and parietal suture between LM3 and 27 (left) | 3 |
| Curve14 | Parietal and squamosal suture between LM26 and 32 (right) | 6 |
| Curve15 | Parietal and squamosal suture between LM27 and 33 (right) | 6 |
| Curve16 | Parietal and interparietal suture between LM34 and 4 (right) | 2 |
| Curve17 | Parietal and interparietal suture between LM35 and 4 (right) | 2 |
| Curve18 | Interparietal and occipital suture between LM34 and 5 (right) | 3 |
| Curve19 | Interparietal and occipital suture between LM35 and 5 (right) | 3 |
| Curve20 | Lateral most edge of occipital between LM34 and 36 (right) | 3 |
| Curve21 | Lateral most edge of occipital between LM35 and 37 (right) | 3 |
| Curve22 | Dorsal and posterior most edge of foramen magnum between LM6 and 36 (right) | 2 |
| Curve23 | Dorsal and posterior most edge of foramen magnum between LM6 and 37 (left) | 2 |
| Curve24 | Ventral and posterior most edge of foramen magnum between LM36 and 7 (right) | 3 |
| Curve25 | Ventral and posterior most edge of foramen magnum between LM37 and 7 (left) | 3 |
| Curve26 | Posterior outline of auditory bulla between LM38 and LM40 (right) | 5 |
| Curve27 | Posterior outline of auditory bulla between LM39 and LM41 (left) | 5 |
| Curve28 | Anterior outline of auditory bulla between LM40 and LM54 (right) | 3 |
| Curve29 | Anterior outline of auditory bulla between LM41 and LM55 (left) | 3 |
| Curve30 | Ventral surface of pterygoid between LM42 and LM 44 (right) | 2 |
| Curve31 | Ventral surface of pterygoid between LM43 and LM 45 (left) | 2 |
| Curve32 | Lingual edge of tooth row between LM44 and 46 (right) | 3 |
| Curve33 | Lingual edge of tooth row between LM45 and 47 (right) | 3 |
| Curve34 | Posterior most edge of maxilla between LM46 and 48 (right) | 3 |
| Curve35 | Posterior most edge of maxilla between LM47 and 49 (left) | 3 |
| Curve36 | Lateral most edge of maxilla between LM48 and 20 (right) | 2 |
| Curve37 | Lateral most edge of maxilla between LM49 and 21 (left) | 2 |
| Curve38 | Anterior edge of lateral supraorbital between LM58 and 20 (right) | 2 |
| Curve39 | Anterior edge of lateral supraorbital between LM59 and 21 (left) | 2 |
| Curve40 | Lateral and anterior edge of orbit between LM20 and 22 (right) | 1 |
| Curve41 | Lateral and anterior edge of orbit between LM21 and 23 (right) | 1 |
| Curve42 | Medial edge of supraorbital between LM22 and 18 (right) | 3 |
| Curve43 | Medial edge of supraorbital between LM22 and 18 (left) | 3 |
| Curve44 | Inciscor root and ventral supraorbital between LM18 and 58 (right) | 3 |
| Curve45 | Inciscor root and ventral supraorbital between LM19 and 59 (left) | 3 |
| Curve46 | Anterior edge of squamosal between LM26 and 28 (right) | 3 |
| Curve47 | Anterior edge of squamosal between LM27 and 29 (left) | 3 |
| Curve48 | Intersection of squamosal with the zygomatic arch between LM28 and 30 (right) | 2 |
| Curve49 | Intersection of squamosal with the zygomatic arch between LM29 and 31 (left) | 2 |
| Patch1 | Nasal surface between Curve1 and Curve 8 (right) | 5 |
| Patch2 | Nasal surface between Curve1 and Curve9 (left) | 5 |
| Patch3 | Frontal surface between Curve2, Curve 10, and Curve12 (right) | 7 |
| Patch4 | Frontal surface between Curve2, Curve 11, and Curve13 (left) | 7 |
| Patch5 | Parietal surface between Curve3, Curve12, Curve14, and 16 (right) | 14 |
| Patch6 | Parietal surface between Curve3, Curve13, 15 and 17 (left) | 14 |
| Patch7 | Interparietal surface between Curve4, Curve16, and Curve18 (right) | 3 |
| Patch8 | Interparietal surface between Curve4, Curve17, and Curve19 (left) | 3 |
| Patch9 | Occipital surface between Curve5, Curve16, Curve20, and Curve22 (right) | 8 |
| Patch10 | Occipital surface between Curve5, Curve17, Curve21, and Curve23 (left) | 8 |
| Patch11 | Maxillar surface between Curve34, Curve36, and Curve38 (right) | 8 |
| Patch12 | Maxillar surface between Curve35, Curve37, and Curve39 (left) | 8 |
| Patch13 | Auditory bullae between Curve28 and 30 (right) | 10 |
| Patch14 | Auditory bullae between Curve29 and 31 (left) | 10 |
| Patch15 | Squamosal surface between Curve14 and 48 (right) | 7 |
| Patch16 | Squamosal surface between Curve15 and 49 (left) | 7 |
|  | **Fixed Landmark subtotal** | **60** |
|  | **Curve Semilandmark subtotal** | **141** |
|  | **Patch Semilandmark subtotal** | **124** |
|  | **GRAND TOTAL** | **325** |
