## Supplementary material for "Australian rodents reveal conserved Cranial Evolutionary Allometry across 10 million years of murid evolution": table A3

**Table A3:** additional alignments of species added to those of Smissen and Rowe (2018) to increase coverage of Australian taxa.

**Additional Alignments**

*Rattus tunneyi*

*Rattus sordidus*

*Rattus lutreolus*

*Rattus fuscipes*

*Melomys burtoni*

*Melomys capensis*
