## Supplementary material for "Australian rodents reveal conserved Cranial Evolutionary Allometry across 10 million years of murid evolution": table A4

**Table A4:** homogeneity of slopes test. P-values for pairwise comparisons of species-level static allometric slopes. Significant p-values (alpha = 0.01) are highlighted in red.

| | **H chr** | **L con** | **L for** | **M bur** | **M cap** | **M cer** | **M fus** | **M gou** | **M mus** | **N ale** | **N cer** | **N fus** | **N mit** | **P apo** | **P aus** | **P del** | **P des** | **P gra** |
|---|---|---|---|---|---|---|---|---|---|---|---|---|---|---|---|---|---|---|
| **C pen** | 0.192 | 0.172 | 0.375 | 0.679 | 0.511 | 0.551 | 0.551 | 0.224 | 0.307 | 0.629 | 0.204 | 0.469 | 0.447 | 0.19 | 0.493 | 0.359 | 0.822 | 0.549 |
| **H chr** | 1 | 0.022 | 0.03 | 0.098 | 0.036 | 0.04 | 0.315 | 0.11 | 0.072 | 0.152 | 0.046 | 0.244 | 0.112 | 0.014 | 0.162 | 0.291 | 0.373 | 0.114 |
| **L con** | | 1 | 0.094 | 0.244 | 0.05 | 0.02 | 0.333 | 0.028 | 0.026 | 0.016 | 0.04 | 0.16 | 0.144 | 0.03 | 0.168 | 0.521 | 0.277 | 0.026 |
| **L for** | | | 1 | 0.064 | 0.07 | 0.082 | 0.068 | 0.194 | 0.092 | 0.439 | 0.076 | 0.277 | 0.253 | 0.232 | 0.058 | 0.044 | 0.12 | 0.19 |
| **M bur** | | | | 1 | 0.285 | 0.493 | 0.21 | 0.347 | 0.024 | 0.623 | 0.12 | 0.383 | 0.914 | 0.064 | 0.457 | 0.158 | 0.651 | 0.749 |
| **M cap** | | | | | 1 | 0.964 | 0.463 | 0.633 | 0.032 | 0.305 | 0.132 | 0.058 | 0.856 | 0.052 | 0.691 | 0.136 | 0.737 | 0.283 |
| **M cer** | | | | | | 1 | 0.459 | 0.431 | 0.072 | 0.261 | 0.072 | 0.088 | 0.76 | 0.028 | 0.747 | 0.255 | 0.653 | 0.359 |
| **M fus** | | | | | | | 1 | 0.776 | 0.012 | 0.487 | 0.192 | 0.086 | 0.681 | 0.024 | 0.186 | 0.202 | 0.679 | 0.144 |
| **M gou** | | | | | | | | 1 | 0.098 | 0.148 | 0.152 | 0.062 | 0.291 | 0.224 | 0.487 | 0.152 | 0.884 | 0.525 |
| **M mus** | | | | | | | | | 1 | 0.158 | 0.012 | 0.2 | 0.088 | 0.032 | 0.064 | 0.062 | 0.022 | 0.088 |
| **N ale** | | | | | | | | | | 1 | 0.248 | 0.05 | 0.449 | 0.2 | 0.22 | 0.224 | 0.697 | 0.517 |
| **N cer** | | | | | | | | | | | 1 | 0.03 | 0.202 | 0.176 | 0.058 | 0.22 | 0.669 | 0.084 |
| **N fus** | | | | | | | | | | | | 1 | 0.024 | 0.03 | 0.297 | 0.355 | 0.481 | 0.214 |
| **N mit** | | | | | | | | | | | | | 1 | 0.172 | 0.259 | 0.24 | 0.667 | 0.778 |
| **P apo** | | | | | | | | | | | | | | 1 | 0.022 | 0.01 | 0.186 | 0.122 |
| **P aus** | | | | | | | | | | | | | | | 1 | 0.09 | 0.503 | 0.042 |
| **P del** | | | | | | | | | | | | | | | | 1 | 0.08 | 0.118 |
| **P des** | | | | | | | | | | | | | | | | | 1 | 0.774 |
| **P gra** | | | | | | | | | | | | | | | | | | 1 |
| **P her** | | | | | | | | | | | | | | | | | | |
| **P hig** | | | | | | | | | | | | | | | | | | |
| **P mol** | | | | | | | | | | | | | | | | | | |
| **P nov** | | | | | | | | | | | | | | | | | | |
| **P ora** | | | | | | | | | | | | | | | | | | |
| **P pat** | | | | | | | | | | | | | | | | | | |
| **P sho** | | | | | | | | | | | | | | | | | | |

| P her | P hig | P mol | P nov | P ora | P pat | P sho | R fus | R leu | R lut | R nor | R rat | R sor | R tun | R vil | U cau | X myo | Z arg | Z ped |
|---|---|---|---|---|---|---|---|---|---|---|---|---|---|---|---|---|---|---|
| 0.417 | 0.569 | 0.461 | 0.084 | 0.569 | 0.749 | 0.529 | 0.479 | 0.483 | 0.633 | 0.619 | 0.749 | 0.541 | 0.561 | 0.467 | 0.84 | 0.238 | 0.752 | 0.487 |
| 0.182 | 0.06 | 0.024 | 0.01 | 0.086 | 0.445 | 0.064 | 0.048 | 0.092 | 0.026 | 0.022 | 0.078 | 0.046 | 0.078 | 0.044 | 0.888 | 0.277 | 0.086 | 0.034 |
| 0.194 | 0.044 | 0.066 | 0.04 | 0.094 | 0.198 | 0.042 | 0.188 | 0.098 | 0.058 | 0.128 | 0.257 | 0.07 | 0.08 | 0.13 | 0.93 | 0.176 | 0.046 | 0.216 |
| 0.006 | 0.18 | 0.248 | 0.01 | 0.255 | 0.084 | 0.236 | 0.303 | 0.008 | 0.024 | 0.585 | 0.172 | 0.042 | 0.124 | 0.086 | 0.8 | 0.01 | 0.02 | 0.182 |
| 0.114 | 0.599 | 0.529 | 0.01 | 0.852 | 0.375 | 0.643 | 0.731 | 0.192 | 0.78 | 0.988 | 0.852 | 0.912 | 0.82 | 0.938 | 0.946 | 0.076 | 0.341 | 0.697 |
| 0.184 | 0.331 | 0.719 | 0.014 | 0.543 | 0.236 | 0.98 | 0.976 | 0.138 | 0.421 | 0.998 | 0.814 | 0.585 | 0.164 | 0.263 | 0.908 | 0.078 | 0.75 | 0.836 |
| 0.206 | 0.19 | 0.671 | 0.024 | 0.409 | 0.22 | 0.982 | 0.996 | 0.118 | 0.339 | 0.976 | 0.77 | 0.481 | 0.062 | 0.285 | 0.906 | 0.072 | 0.888 | 0.89 |
| 0.202 | 0.75 | 0.311 | 0.042 | 0.299 | 0.23 | 0.333 | 0.615 | 0.425 | 0.677 | 0.758 | 0.699 | 0.575 | 0.375 | 0.212 | 0.842 | 0.046 | 0.331 | 0.363 |
| 0.439 | 0.246 | 0.457 | 0.291 | 0.437 | 0.455 | 0.375 | 0.224 | 0.246 | 0.309 | 0.116 | 0.096 | 0.186 | 0.152 | 0.118 | 0.856 | 0.437 | 0.621 | 0.208 |
| 0.052 | 0.066 | 0.06 | 0.014 | 0.08 | 0.098 | 0.056 | 0.204 | 0.062 | 0.024 | 0.138 | 0.144 | 0.028 | 0.032 | 0.052 | 0.653 | 0.008 | 0.1 | 0.192 |
| 0.13 | 0.25 | 0.178 | 0.008 | 0.489 | 0.469 | 0.367 | 0.281 | 0.032 | 0.146 | 0.938 | 0.523 | 0.265 | 0.397 | 0.144 | 0.886 | 0.136 | 0.222 | 0.138 |
| 0.088 | 0.112 | 0.068 | 0.004 | 0.068 | 0.379 | 0.06 | 0.04 | 0.052 | 0.106 | 0.09 | 0.138 | 0.118 | 0.072 | 0.032 | 0.822 | 0.232 | 0.076 | 0.09 |
| 0.084 | 0.068 | 0.493 | 0.016 | 0.06 | 0.443 | 0.06 | 0.056 | 0.034 | 0.05 | 0.236 | 0.467 | 0.305 | 0.355 | 0.663 | 0.966 | 0.792 | 0.014 | 0.082 |
| 0.158 | 0.617 | 0.629 | 0.06 | 0.597 | 0.321 | 0.936 | 0.737 | 0.104 | 0.579 | 0.98 | 0.75 | 0.651 | 0.271 | 0.355 | 0.866 | 0.124 | 0.563 | 0.766 |
| 0.02 | 0.05 | 0.022 | 0.13 | 0.158 | 0.24 | 0.088 | 0.056 | 0.018 | 0.054 | 0.2 | 0.032 | 0.048 | 0.058 | 0.038 | 0.601 | 0.022 | 0.042 | 0.032 |
| 0.118 | 0.337 | 0.914 | 0.004 | 0.202 | 0.313 | 0.541 | 0.952 | 0.176 | 0.218 | 0.996 | 0.93 | 0.643 | 0.393 | 0.691 | 0.952 | 0.096 | 0.138 | 0.423 |
| 0.058 | 0.234 | 0.257 | 0.002 | 0.126 | 0.018 | 0.2 | 0.343 | 0.349 | 0.325 | 0.363 | 0.541 | 0.373 | 0.297 | 0.403 | 0.78 | 0.038 | 0.148 | 0.467 |
| 0.168 | 0.595 | 0.633 | 0.068 | 0.657 | 0.842 | 0.603 | 0.575 | 0.623 | 0.778 | 0.852 | 0.633 | 0.695 | 0.607 | 0.545 | 0.948 | 0.136 | 0.633 | 0.483 |
| 0.082 | 0.311 | 0.545 | 0.022 | 0.824 | 0.429 | 0.627 | 0.619 | 0.012 | 0.226 | 0.966 | 0.727 | 0.309 | 0.098 | 0.435 | 0.894 | 0.066 | 0.102 | 0.351 |
| 1 | 0.222 | 0.128 | 0.016 | 0.15 | 0.146 | 0.14 | 0.287 | 0.515 | 0.293 | 0.333 | 0.405 | 0.192 | 0.14 | 0.164 | 0.816 | 0.01 | 0.307 | 0.25 |
| | 1 | 0.096 | 0.068 | 0.806 | 0.329 | 0.457 | 0.549 | 0.03 | 0.447 | 0.804 | 0.475 | 0.232 | 0.142 | 0.13 | 0.84 | 0.064 | 0.361 | 0.106 |
| | | 1 | 0.034 | 0.347 | 0.168 | 0.711 | 0.97 | 0.132 | 0.15 | 0.808 | 0.794 | 0.545 | 0.305 | 0.657 | 0.938 | 0.142 | 0.547 | 0.665 |
| | | | 1 | 0.07 | 0.05 | 0.036 | 0.056 | 0.026 | 0.038 | 0.03 | 0.02 | 0.016 | 0.008 | 0.014 | 0.291 | 0.004 | 0.026 | 0.078 |
| | | | | 1 | 0.443 | 0.8 | 0.497 | 0.028 | 0.523 | 0.872 | 0.587 | 0.451 | 0.349 | 0.477 | 0.874 | 0.054 | 0.321 | 0.371 |
| | | | | | 1 | 0.265 | 0.271 | 0.214 | 0.355 | 0.393 | 0.389 | 0.355 | 0.405 | 0.345 | 0.938 | 0.263 | 0.23 | 0.18 |
| | | | | | | 1 | 0.986 | 0.166 | 0.575 | 0.996 | 0.94 | 0.81 | 0.257 | 0.553 | 0.904 | 0.036 | 0.974 | 0.946 |

| | | | | | | | | | | | | |
|---|---|---|---|---|---|---|---|---|---|---|---|---|
| **R fus** | 1 | 0.255 | 0.375 | 0.77 | 0.796 | 0.511 | 0.16 | 0.371 | 0.912 | 0.136 | 0.988 | 1 |
| **R leu** | | 1 | 0.423 | 0.112 | 0.693 | 0.427 | 0.04 | 0.152 | 0.942 | 0.118 | 0.301 | 0.557 |
| **R lut** | | | 1 | 0.353 | 0.725 | 0.904 | 0.403 | 0.23 | 0.944 | 0.058 | 0.693 | 0.794 |
| **R nor** | | | | 1 | 0.615 | 0.681 | 0.719 | 0.325 | 0.896 | 0.078 | 0.984 | 0.754 |
| **R rat** | | | | | 1 | 0.984 | 0.91 | 0.884 | 0.958 | 0.226 | 0.776 | 0.794 |
| **R sor** | | | | | | 1 | 0.916 | 0.591 | 0.966 | 0.136 | 0.483 | 0.723 |
| **R tun** | | | | | | | 1 | 0.888 | 0.976 | 0.134 | 0.096 | 0.198 |
| **R vil** | | | | | | | | 1 | 0.982 | 0.174 | 0.269 | 0.561 |
| **U cau** | | | | | | | | | 1 | 0.984 | 0.9 | 0.898 |
| **X myo** | | | | | | | | | | 1 | 0.02 | 0.074 |
| **Z arg** | | | | | | | | | | | 1 | 0.86 |
| **Z ped** | | | | | | | | | | | | 1 |
