## Supplementary material for "Australian rodents reveal conserved Cranial Evolutionary Allometry across 10 million years of murid evolution": table A5

**Table A5:** phylogenetic Procrustes ANCOVA. An alternative analysis to table 1*A* that uses mean shapes for each species and considers the phylogenetic tree.

|  | **Df** | **SS** | **MS** | **Rsq** | **F** | **Z** | **Pr(>F)** |
| --- | --- | --- | --- | --- | --- | --- | --- |
| **log(size)** | 1 | 0.016 | 0.016 | 0.407 | 25.148 | 5.355 | 0.002 |
| **clade** | 7 | 0.003 | 0 | 0.087 | 0.771 | -1.063 | 0.856 |
| **log(size):clade** | 5 | 0.005 | 0.001 | 0.134 | 1.656 | 2.222 | 0.02 |
| **Residuals** | 23 | 0.015 | 0.001 | 0.372 |  |  |  |
| **Total** | 36 | 0.039 |  |  |  |  |  |
