## Supplementary material for "Australian rodents reveal conserved Cranial Evolutionary Allometry across 10 million years of murid evolution": table A6

**Table A6:** comparative evolutionary rates. Upper triangle reports relative rates between groups (row to column) and the lower triangle reports the p-value (corrected for multiple comparisons using the Bonferroni method). Species were assigned to groups as defined in the Methods and in figure 4*B*.

|  | **Non-specialist** | **Frugivore** | **Folivore** | **Carnivore** | **Hopping** |
| --- | --- | --- | --- | --- | --- |
| **Non-specialist** | - | 3.96 | 2.48 | 2.13 | 1.06 |
| **Frugivore** | 0.04 | - | 1.6 | 1.86 | 4.19 |
| **Folivore** | 0.1 | 1 | - | 1.16 | 2.63 |
| **Carnivore** | 1 | 1 | 1 | - | 2.26 |
| **Hopping** | 1 | 0.08 | 0.14 | 1 | - |
