## Supplementary material for "Australian rodents reveal conserved Cranial Evolutionary Allometry across 10 million years of murid evolution": fig. A1

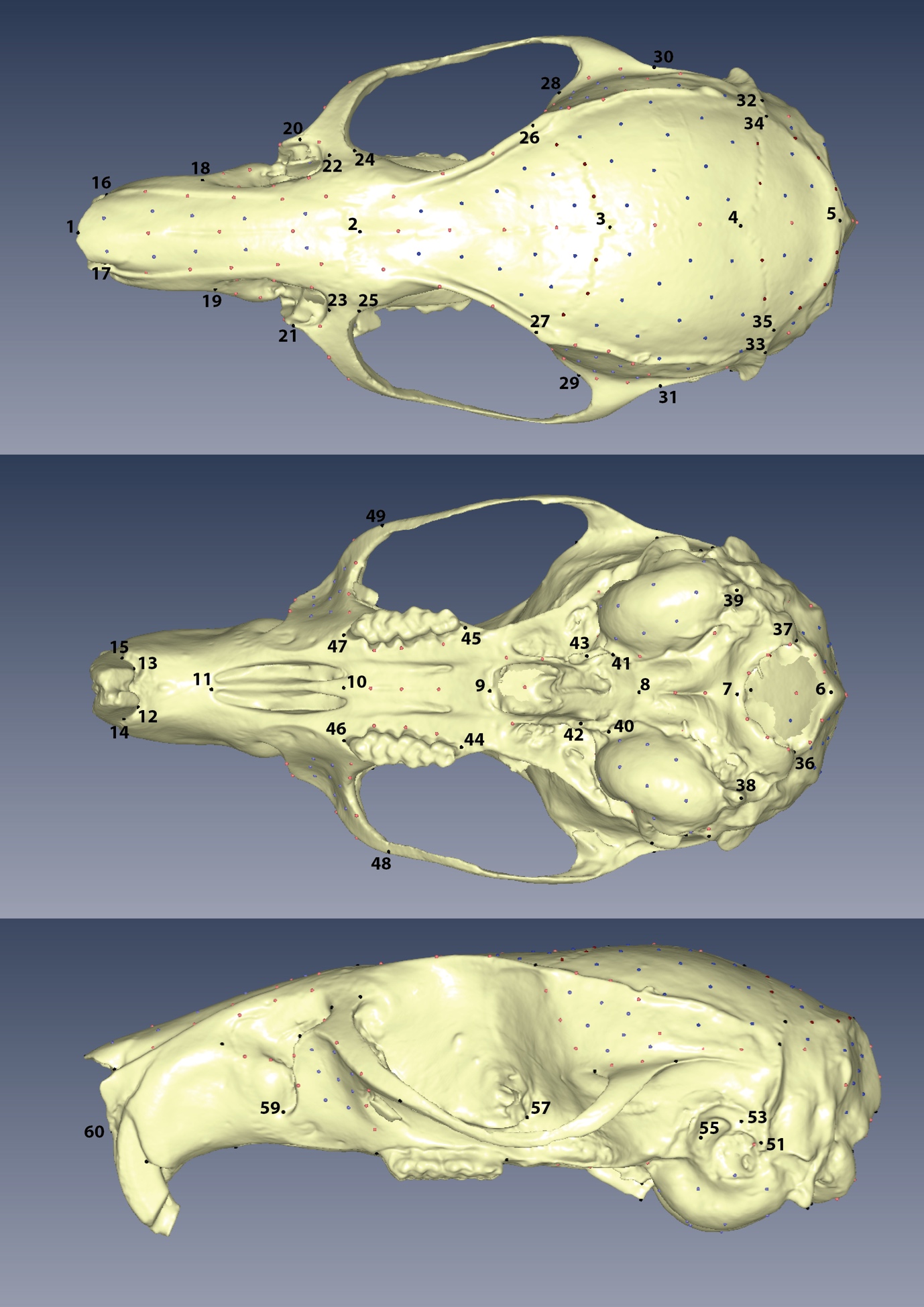

**Figure A1:** placement of landmarks in our protocol on a typical cranium. See table A1 for definitions of each landmark, semi-landmark curve, and patch semi-landmark.
