## Supplementary material for "Australian rodents reveal conserved Cranial Evolutionary Allometry across 10 million years of murid evolution": fig. A2

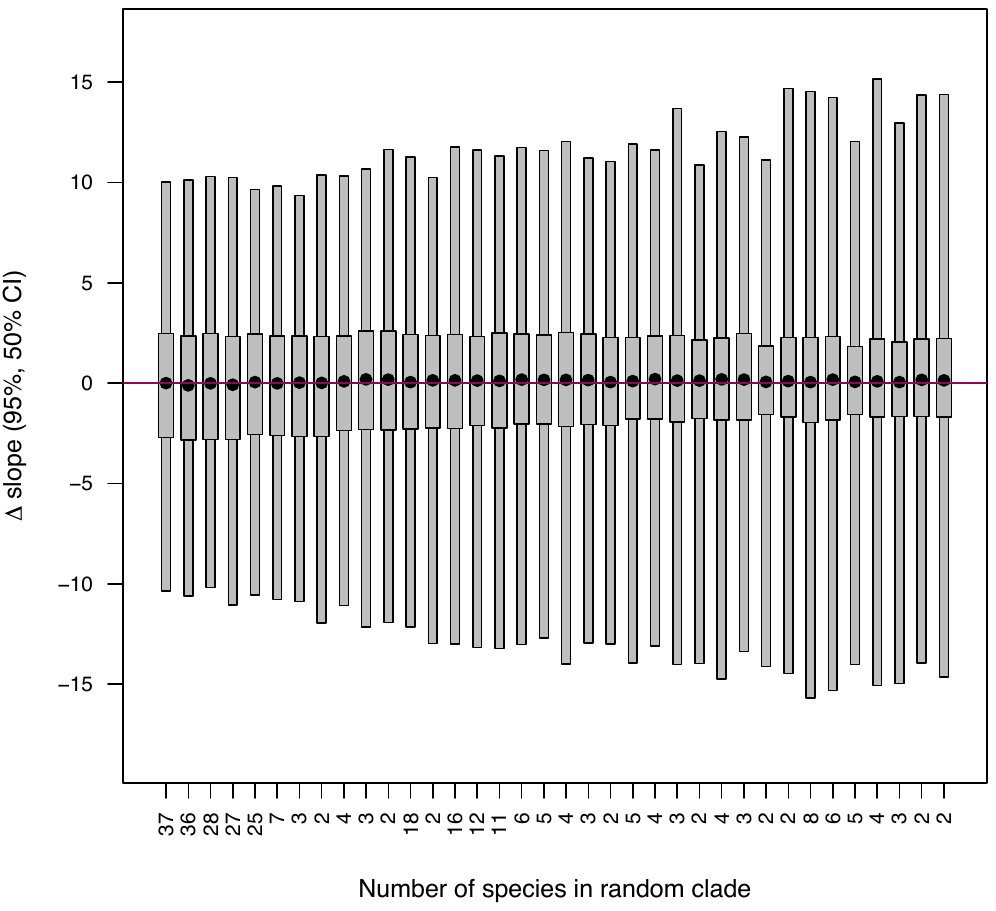

**Figure A2:** boxplot showing confidence intervals (95%, 50%) and the median value (black point) for each of the clades after 100 replicates with randomized clades, i.e. species could be randomized from across the phylogenetic tree. Plot is comparable to figure 3*A* with clades ordered in the exact same size bins, but single species bins ignored.
